# Simultaneous *in vivo* time-lapse stiffness mapping and fluorescence imaging of developing tissue

**DOI:** 10.1101/323501

**Authors:** Amelia J. Thompson, Iva K. Pillai, Ivan B. Dimov, Christine E. Holt, Kristian Franze

## Abstract

Tissue mechanics is important for development; however, the spatio-temporal dynamics of *in vivo* tissue stiffness is still poorly understood. We here developed tiv-AFM, combining time-lapse *in vivo* atomic force microscopy with upright fluorescence imaging of embryonic tissue, to show that in the developing *Xenopus* brain, a stiffness gradient evolves over time because of differential cell proliferation. Subsequently, axons turn to follow this gradient, underpinning the importance of time-resolved mechanics measurements.

---

During embryonic development, many biological processes are regulated by tissue mechanics *in vivo*, including cell migration^1^, neuronal growth^2^, and large-scale tissue remodelling^3,4^. Recent measurements at specific time points suggested that tissue mechanics change during developmental^2,5,6^ and pathological^7,8^ processes, which might significantly impact cell function. Furthermore, several approaches have recently been developed to measure *in vivo* tissue stiffness, including atomic force microscopy^1,2,9^, magnetic resonance elastography^10^, Brillouin microscopy^11^, and magnetically responsive ferrofluid microdroplets^12^. However, the precise spatiotemporal dynamics of tissue mechanics remains poorly understood, and how cells respond to changes in local tissue stiffness *in vivo* largely unknown.

To enable time-resolved measurements of developmental tissue mechanics, we here developed time-lapse *in vivo* atomic force microscopy (tiv-AFM), a method that combines sensitive upright epifluorescence imaging of opaque samples such as frog embryos with iterated AFM indentation measurements of *in vivo* tissue at cellular resolution and at a time scale of minutes (Fig. 1). A fluorescence zoom stereomicroscope equipped with an sCMOS camera (quantum yield 82%) were custom-fitted above a bio-AFM set-up (Supplementary Fig. 1), which had a transparent pathway along the area of the cantilever. To cope with the long working distances required for imaging through the AFM head, the microscope was fitted with a 0.125 NA / 114 mm WD objective. The AFM was set up on an automated motorised stage containing a temperature-controlled sample holder to maintain live specimens at optimal conditions during the experimental time course. (Fig. 1a, b) (see online methods for details).

**Figure 1.**
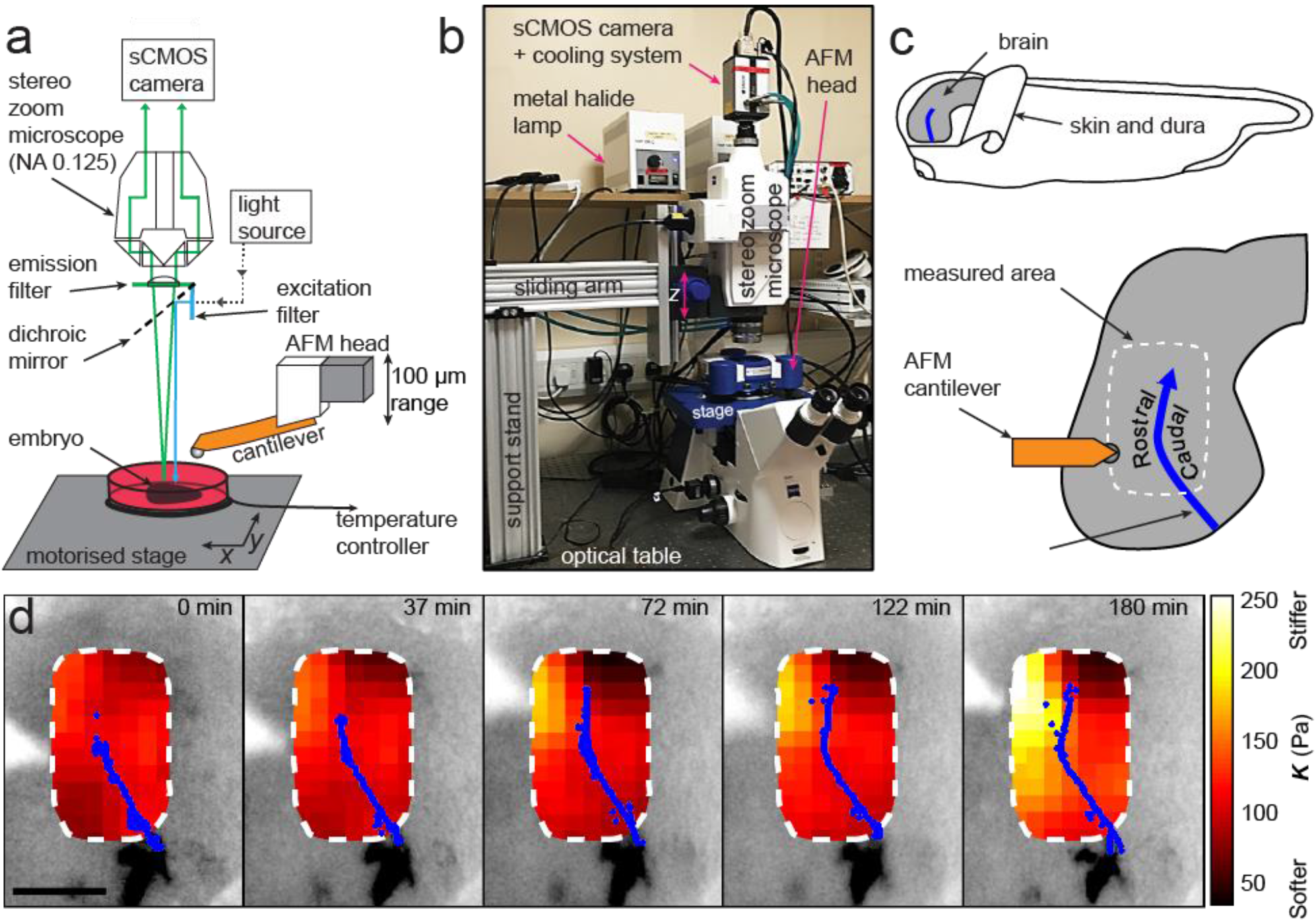
Combined time-lapse *in vivo* AFM (tiv-AFM). (a) Schematic (not to scale) and (b) photograph of the experimental setup. An AFM with 100 μm *z*-piezo range is positioned above a temperature-controlled sample chamber containing the specimen. A custom-fit fluorescence zoom stereomicroscope with a long (114 mm) working distance and NA 0.125 objective, connected to a high quantum-efficiency sCMOS camera, is mounted on a custom-built support stand above the AFM head optimised for trans-illumination. The specimen is moved by a motorised *x/y* stage to allow AFM-based mapping of large areas. (c) Schematic of a*Xenopus* embryo, showing how the brain is prepared for tiv-AFM (top) and close-up of the brain (bottom) showing the approximate region measured, within which optic tract (OT) axons (blue) turn caudally. (d) Time-lapse stiffness maps obtained from a tiv-AFM experiment, showing outlines of fluorescently labelled OT axons (blue) and processed AFM-based stiffness maps (colour maps) overlaid on images of the brain. Colour maps encode the apparent elastic modulus *K*, a measure of tissue stiffness, assessed at an indentation force *F* = 10 nN. The time in minutes on each frame is taken from the timestamp of the first measurement in each successive stiffness map; the corresponding overlaid fluorescence images were obtained simultaneously. AFM measurement resolution, 20 μm; scale bar, 100 μm.

We tested tiv-AFM using the developing *Xenopus* embryo brain during outgrowth of the optic tract (OT) as a model (Fig. 1c). In the OT, retinal ganglion cell axons grow in a bundle across the brain surface, making a stereotypical turn in the caudal direction *en route* that directs them to their target, the visual centre of the brain^13^. We previously demonstrated that by later stages of OT outgrowth (i.e. when axons had reached their target), a local stiffness gradient lies orthogonal to the path of OT axons, with the stiffer region rostral to the OT and softer region caudal to it^2^. This gradient strongly correlated with axon turning, with the OT routinely turning caudally towards softer tissue^2^. We therefore wanted to determine when this stiffness gradient first developed, whether its emergence preceded OT axon turning, and what the origin of that stiffness gradient was.

To answer these questions, we performed iterated tiv-AFM measurements of the embryonic brain *in vivo* at early-intermediate stages, i.e. just before and during turn initiation by the first ‘pioneer’ OT axons. The apparent elastic modulus *K*, which is a measure of the tissue’s elastic stiffness, was assessed in a ~150 μm by 250 μm raster at 20 μm resolution every ~35 minutes, producing a sequence of ‘stiffness maps’ of the area (Supplementary Fig. 2). To reduce noise, raw AFM data were interpolated and smoothed in *x-,* y-, and time dimensions using an algorithm based on the discrete cosine transform (Fig. 1d, see online methods for details)^14,15^. Simultaneously, we recorded optical time-lapse images of fluorescently labelled retinal ganglion cell axons growing through the region of interest (Fig. 1d, Supplementary Fig. 2a).

Early in the time-lapse sequence (i.e. prior to axon turning), the stiffness of the brain was similar on both sides of the OT. However, over the time course of the measurements a stiffness gradient arose, mostly due to rising stiffness of tissue rostral to the OT (Fig. 1d). Visual inspection of the fold-change in tissue stiffness from one time point to the next indicated that significant changes in tissue mechanics were already occurring approximately 40 – 80 minutes after the onset of measurements (Supplementary Fig. 2c), i.e., before axons started turning caudally, suggesting that the tissue stiffness gradient was established prior to axon turning.

To test this hypothesis, we quantified the temporal evolution of the stiffness gradient in a small region immediately in front of the advancing OT. At the beginning of each time point in the sequence of tiv-AFM maps, we calculated the size of the angle through which axons turned (‘OT turn angle’). For each animal, minimum and maximum absolute values were rescaled to 0 and 1, respectively. The projected appearance of the stiffness gradient significantly preceded the projected onset of axon turning (Fig. 2b), indicating that axons turned after the stiffness gradient was established (Fig. 2a), which is consistent with a role for mechanical gradients in helping to guide OT axons caudally^2^. In line with this idea, RGC axons from heterochronic eye primordia transplants growing through *Xenopus* brains at stages before the stiffness gradient is established grow rather straight and do not turn caudally in the mid-diencephalon^16^.

**Figure 2.**
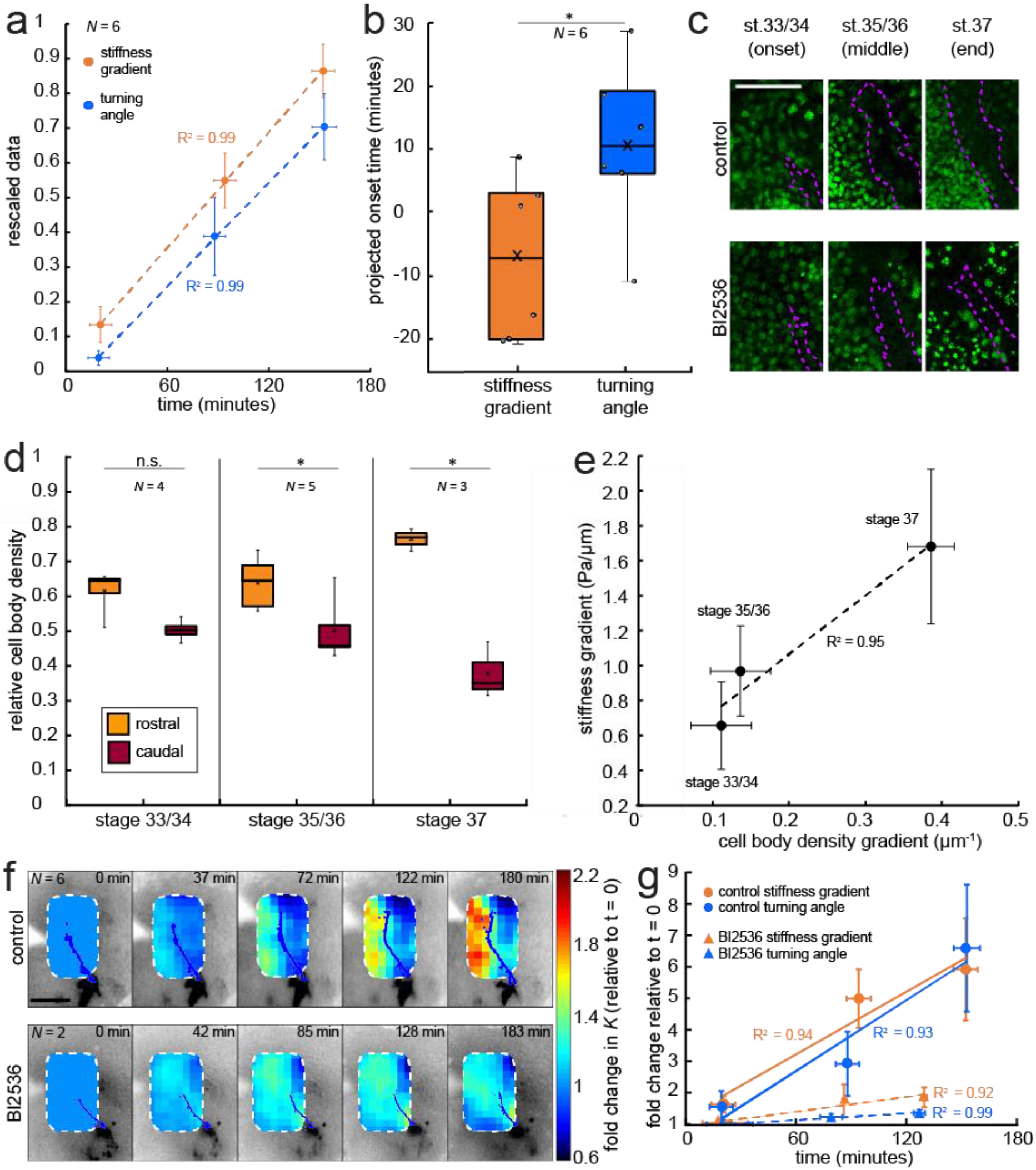
Development of mechanical gradients in *Xenopus* embryo brains precedes axon turning and is underpinned by changes in local cell body densities. (a) Plot of mean re-scaled values for the stiffness gradient (orange) and OT turn angle (blue). Dashed lines denote linear fits (R^2^ = 0.99). (b) Boxplots of the extrapolated appearance times of the stiffness gradients and the onset of OT axon turning, relative to the start time of tiv-AFM measurements. Extrapolations are based on linear fits to the re-scaled data for individual animals. Stiffness gradients appear significantly earlier than the onset of axon turning (*p* = 0.03, paired Wilcoxon signed-rank test). (c) Immunohistochemistry of nuclei (green) in whole-mount control (middle row) and mitotic inhibitor-treated (bottom) *Xenopus* embryo brains at successive developmental stages. Stages shown correspond to the onset (stage 33/34), middle (stage 35/36), and end (stage 37) of tiv-AFM measurements. OT axons are outlined in purple. (d) Local cell body densities were significantly higher rostral to the OT than caudal to it at both stage 35/36 (*p* = 0.03, paired Wilcoxon signed-rank test) and stage 37 (*p* = 0.04). (e) Correlation between gradients in local cell body density and tissue stiffness. Binned absolute values for the stiffness gradient (in Pa / μm) are plotted against the mean cell density gradient at each developmental stage. Dashed line denotes linear fit (R^2^ = 0.95; Pearson’s correlation coefficient *p* = 0.97). (f) Time-lapse AFM montages showing fold-changes in brain stiffness in representative control (top; OT axons outlined in blue) and mitotic inhibitor-treated embryos (bottom; OT axons in magenta). The colour scale encodes the fold-change in *K* at each location on the stiffness map, expressed relative to the values obtained at t = 0 minutes. (g) Plots of the fold-change over time in stiffness gradient (orange) and OT turn angle (blue) for both control and mitotic inhibitor-treated embryos. Blocking mitosis significantly attenuated the rise in both stiffness gradient (*p* = 0.01, linear regression analysis) and OT axon turning (*p* = 0.01). Boxplots show median, 1^st^, and 3^rd^ quartiles; whiskers show the spread of the data; ‘×’ indicates the mean. Error bars denote standard error of the mean. \**p* < 0.05. AFM measurement resolution, 20 μm; scale bars, 100 μm. *N* denotes number of animals.

We have previously shown that tissue stiffness scales with local cell body density^1,17^, and that in *Xenopus* embryo brains local stiffness gradients at later developmental stages (39/40) correlate with a gradient in cell density^2^. To determine if changes in cell densities are driving changes in tissue stiffness, and thus parallel the evolution of the stiffness gradient at earlier stages, we assayed cell densities using DAPI labelling of nuclei in whole-mounted brains with fluorescently labelled OTs, beginning at the morphological stage corresponding to the start of tiv-AFM measurements (33/34) and repeated for the two subsequent stages encompassing OT turning (35/36 and 37).

While at the first stage cell densities on both sides of the OT were similar, a clear difference in nuclear densities rostral and caudal to the OT developed at later stages (Fig. 2c). Cell densities at the two later stages were significantly higher in the region rostral to the OT (i.e. where tissue was stiffer) than caudal to it, and the overall magnitude of the cell density gradient significantly rose over time (Fig. 2d). Plotting the stage-specific gradient in cell body densities against the stiffness gradient revealed a strong linear correlation between them (Pearson’s correlation coefficient *p* = 0.97) (Fig. 2e).

To test if local cell densities drive the evolution of the stiffness gradient during OT turning, we repeated both nuclear staining and tiv-AFM measurements on embryos treated with the mitotic blocker BI2536^18^, which has previously been used to inhibit *in vivo* cell proliferation in the embryonic retina^19^. In BI2536-treated brains, the nuclear density rostral to the OT and thus the cell density gradient was significantly decreased compared to controls, particularly at later stages (Fig. 2c). Blocking cell proliferation furthermore significantly attenuated both the stiffness gradient and the OT turn angle (Fig. 2f, g), suggesting that the gradient in cell densities is responsible for the stiffness gradient, which on the other hand instructs axon growth.

Tiv-AFM allows simultaneous time-lapse measurements of cell and tissue mechanics *in vivo* and optical monitoring of fluorescently labelled structures at the surface of otherwise optically opaque samples at length and time scales that are relevant for developmental processes. It enabled us for the first time to trace the *in vivo* mechanical properties of the embryonic *Xenopus* brain as the embryo developed, and to relate mechanical tissue changes to a key event in axon pathfinding.

More broadly, tiv-AFM can also be easily adapted for *in vivo* applications in other small organisms, or alternatively in tissues *ex vivo.* It can be used to study cellular responses to a range of mechanical stimuli via the AFM *in vivo,* such as sustained compression^1,2^, or to track the temporal mechanical response of tissues or organs to different pharmacological treatments (such as the mitotic inhibitor used here). Additionally, the setup is very versatile and can be further expanded, for example, by combining it with calcium imaging to investigate how cellular activity is regulated by changes in tissue stiffness during development and pathology. Tiv-AFM will greatly expand the range of bio-AFM experiments possible, allowing for more scope both for a detailed characterisation of *in vivo* tissue mechanics during development and disease progression, and for testing how mechanics shapes cell behaviour and function.

## Acknowledgements

The authors would like to thank Alex Winkel (JPK Instruments) for technical help, as well as the Wellcome Trust (grant to A.J.T.), the Cambridge Philosophical Society and the Cambridge Trusts (studentships to A.J.T.), the Malaysian Commonwealth Studies Centre (funding to E.K.P), the UK Engineering and Physical Sciences Research Council Cambridge NanoDTC (grant EP/L015978/1 to I.B.D.), ERC Advanced Grant 322817 (CEH), the UK Biotechnology and Biological Sciences Research Council (BB/M020630/1 & BB/N006402/1 to K.F.), the UK Medical Research Council (Career Development Award G1100312/1 to K.F.), and the Eunice Kennedy Shriver National Institute Of Child Health & Human Development of the National Institutes of Health under Award Number R21HD080585 (to K.F.) for funding support. The content is solely the responsibility of the authors and does not necessarily represent the official views of the National Institutes of Health.

## Supplementry Figures

**Supplementary Figure 1:**
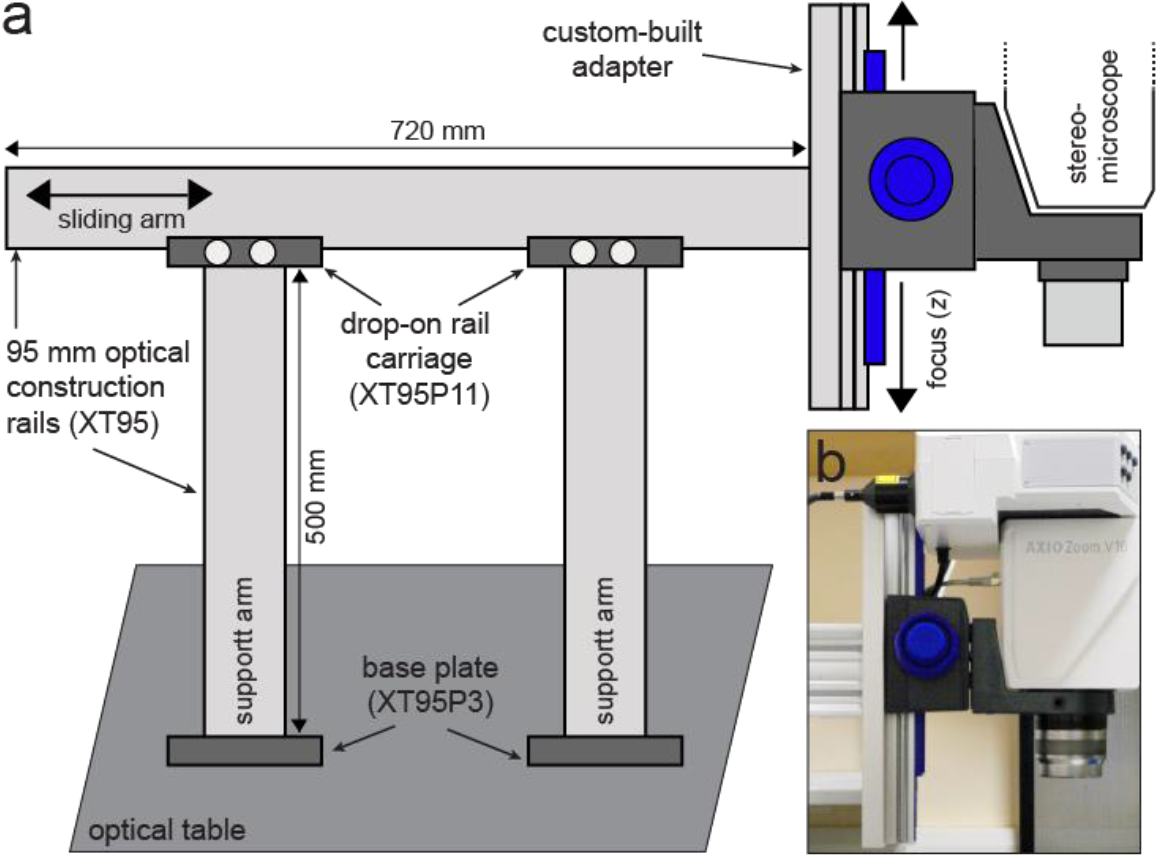
Custom-built support stand for the upright optical imaging set-up. (a) Schematic (not to scale) of support stand and the adapter used to mount the upright fluorescence stereomicroscope to the stand. Codes denote part numbers (Thorlabs) where applicable. (b) Close-up photograph of the custom-built adapter shown in (a), with fluorescence stereomicroscope in place.

**Supplementary Figure 2:**
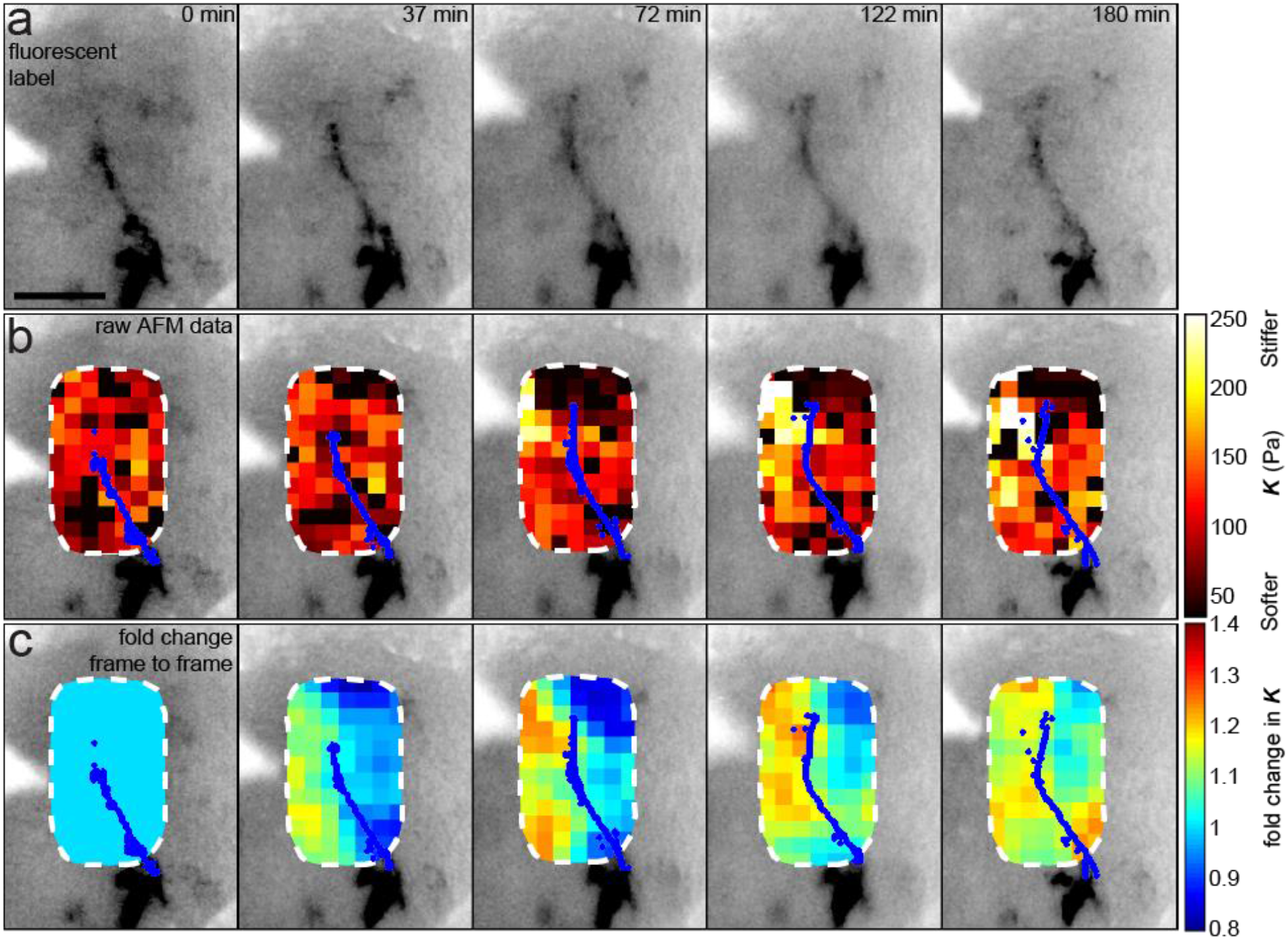
Data processing for tiv-AFM experiments (same *Xenopus* embryo as in Fig. 1d). (a) Inverted epifluorescence raw images of OT axons (black) acquired during tiv-AFM measurements, labelled with membrane-bound GFP under the control of the cell type-specific *ath5* promoter. (b) Overlaid tiv-AFM-based stiffness maps, encoding raw values of apparent elastic moduli *K*, with OT axon outlines shown in blue. Black squares denote points where AFM data were not analysable. To generate the final stiffness maps used for mechanical gradient quantification, missing *K* values were interpolated and data were smoothed in *x*-, *y*-, and *t*- dimensions (*cf*. Fig. 1d; for more details see online methods). (c) Visualisation of fold-changes in brain tissue stiffness from one time point to the next, based on the interpolated and smoothed data shown in Fig. 1d. Colour scale encodes the fold-change in *K* at each location on the stiffness map, expressed relative to the values at the previous time point, with the exception of t = 0 minutes, where all values were set to 1. Tissue stiffness changed throughout the time course, with significant changes already occurring between ~40-80 minutes after the start of the experiment. AFM measurement resolution, 20 μm; scale bar, 100 μm.

## References

1 Barriga, E. H., Franze, K., Charras, G. & Mayor, R. Tissue stiffening coordinates morphogenesis by triggering collective cell migration in vivo. Nature 554, 523–527, (2018).

2 Koser, D. E. et al. Mechanosensing is critical for axon growth in the developing brain. Nat Neurosci 19, 1592–1598, (2016).

3 Butler, L. C. et al. Cell shape changes indicate a role for extrinsic tensile forces in Drosophila germ-band extension. Nat Cell Biol 11, 859–864, (2009).

4 Munjal, A., Philippe, J. M., Munro, E. & Lecuit, T. A self-organized biomechanical network drives shape changes during tissue morphogenesis. Nature 524, 351–355, (2015).

5 Iwashita, M., Kataoka, N., Toida, K. & Kosodo, Y. Systematic profiling of spatiotemporal tissue and cellular stiffness in the developing brain. Development 141, 3793–3798, (2014).

6 Majkut, S. et al. Heart-specific stiffening in early embryos parallels matrix and myosin expression to optimize beating. Curr Biol 23, 2434–2439, (2013).

7 Murphy, M. C. et al. Decreased brain stiffness in Alzheimer’s disease determined by magnetic resonance elastography. J Magn Reson Imaging 34, 494–498, (2011).

8 Moeendarbary, E. et al. The soft mechanical signature of glial scars in the central nervous system. Nat Commun 8, 14787, (2017).

9 Gautier, H. O. et al. Atomic force microscopy-based force measurements on animal cells and tissues. Methods in Cell Biology 125, (2015).

10 Sack, I., Beierbach, B., Hamhaber, U., Klatt, D. & Braun, J. Non-invasive measurement of brain viscoelasticity using magnetic resonance elastography. NMR Biomed 21, 265–271, (2008).

11 Scarcelli, G. & Yun, S. H. In vivo Brillouin optical microscopy of the human eye. Opt Express 20, 9197–9202, (2012).

12 Serwane, F. et al. In vivo quantification of spatially varying mechanical properties in developing tissues. Nat Methods 14, 181–186, (2017).

13 McFarlane, S. & Lom, B. The Xenopus retinal ganglion cell as a model neuron to study the establishment of neuronal connectivity. Developmental Neurobiology 72, 520–536, (2012).

14 Garcia, D. Robust smoothing of gridded data in one and higher dimensions with missing values. Comput Stat Data Anal 54, 1167–1178, (2010).

15 Garcia, D. A fast all-in-one method for automated post-processing of PIV data. Exp Fluids 50, (2011).

16 Cornel, E. & Holt, C. Precocious pathfinding: retinal axons can navigate in an axonless brain. Neuron 9, 1001–1011, (1992).

17 Koser, D. E., Moeendarbary, E., Hanne, J., Kuerten, S. & Franze, K. CNS cell distribution and axon orientation determine local spinal cord mechanical properties. Biophys J 108, 2137–2147, (2015).

18 Lenart, P. et al. The small-molecule inhibitor BI 2536 reveals novel insights into mitotic roles of polo like kinase 1. Curr Biol 17, 304–315, (2007).

19 Weber, I.P. et al. Mitotic position and morphology of committed precursor cells in the zebrafish retina adapt to architectural changes upon tissue maturation. Cell Rep 7, 386–397, (2014).

